# Targeting HLA-E Positive Cancers with a Novel NKG2A/C Switch Receptor

**DOI:** 10.1101/2023.12.14.571754

**Authors:** Michelle Sætersmoen, Ivan S. Kotchetkov, Lamberto Torralba-Raga, Jorge Mansilla-Soto, Ebba Sohlberg, Silje Zandstra Krokeide, Quirin Hammer, Michel Sadelain, Karl-Johan Malmberg

**Affiliations:** Precision Immunotherapy Alliance (PRIMA), University of Oslo, Oslo, Norway; Department of Neurology, Memorial Sloan Kettering Cancer Center, New York, NY, USA; Center for Cell Engineering and Immunology Program, Memorial Sloan Kettering Cancer Center, New York, NY, USA; Department of Cancer Immunology, Institute for Cancer Research, Oslo University Hospital, Oslo, Norway; Departments of Immunology and Bioengineering, H. Lee Moffitt Cancer Center and Research Institute, Tampa, FL, USA; Center for Infectious Medicine, Department of Medicine Huddinge, Karolinska Institutet, Stockholm, Sweden

## Abstract

HLA-E is overexpressed by approximately 80% of solid tumors, including malignant glioblastoma, and is emerging as a major checkpoint for NKG2A^+^ CD8^+^ T cells and NK cells in the tumor microenvironment and circulation. This axis operates side-by-side with PD-L1 to shut down effector responses by T and NK cells. Here, we engineered a novel chimeric A/C switch receptor, combining the strong HLA-E binding affinity of the NKG2A receptor ectodomain with the activating signaling of the NKG2C receptor endodomain. We found that A/C Switch-transduced NK and T cells displayed superior and specific cytotoxic function when challenged with tumor cells exhibiting medium to high HLA-E expression. Furthermore, A/C Switch-expressing human T cells demonstrated enhanced anti-tumor function in a xenograft model of glioblastoma. Importantly, the activity of the modified T cells was governed by an equilibrium between A/C Switch transduction level and HLA-E expression, creating a therapeutic window to safeguard against on-target off-tumor toxicities. Indeed, normal cells remained insensitive to A/C Switch engineered T cells even after pre-treatment with IFN-γ to induce HLA-E expression. We propose that this novel A/C switch receptor may operate alone to control tumor cells expressing high levels of HLA-E or in combination with other engineered specificities to overcome the suppressive NKG2A/HLA-E checkpoint.

## Introduction

Immunotherapy has dramatically shifted treatment paradigms in oncology, but tumor resistance mechanisms continue to hinder responses in many cancer contexts (Cappell and Kochenderfer, 2023; Kalbasi and Ribas, 2020; Kraehenbuehl et al., 2022; Sharma et al., 2023; Sharma et al., 2017). The non-classical HLA class I molecule, HLA-E, has elevated expression in many human tumors (Andersson et al., 2016; Andre et al., 2018; Gooden et al., 2011; Kamiya et al., 2019), including glioblastoma (Hrbac et al., 2022; Mittelbronn et al., 2007; Wischhusen et al., 2005; Wolpert et al., 2012; Wu et al., 2020), and is emerging as a major checkpoint for NKG2A^+^ CD8^+^ T cells and NK cells (Andre et al., 2018; Borst et al., 2022; Herbst et al., 2022; Liu et al., 2023; Salome et al., 2022). Blockade of this suppressive pathway unleashes cytotoxic lymphocytes and can prevent metastasis (Liu et al., 2023). This notion is supported by CRISPR screens in mice, identifying Qa-1 (the mouse orthologue to HLA-E) as a limiting factor for immunotherapy (Dubrot et al., 2022).

HLA-E is recognized by T and NK cells through the CD94/NKG2 family of receptors, which are composed of members with activating or inhibitory potential (Braud et al., 2003; Braud et al., 1998; Houchins et al., 1997; Lee et al., 1998; McMahon and Raulet, 2001; Miller et al., 2003). The CD94/NKG2A heterodimeric receptor transmits inhibitory signals via two immunoreceptor tyrosine-based inhibitory motifs (ITIMs) in the cytoplasmic tail of NKG2A, with downstream events leading to an inhibition of cytokine secretion and cytotoxicity in CD8^+^ T and NK cells (Carretero et al., 1997; Carretero et al., 1998; Le Drean et al., 1998). In contrast, the ligation of CD94/NKG2C by HLA-E transduces an activating signal mediated by the DAP12 adaptor protein bearing an immunoreceptor tyrosine-based activation motif (ITAM) (Lanier, 2009; Lanier et al., 1998). The CD94/NKG2A heterodimer has a six times greater affinity for its ligand than its NKG2C counterpart (Kaiser et al., 2005; Kaiser et al., 2008).

In this study, we engineered a novel chimeric NKG2A/NKG2C (A/C) Switch receptor, combining the strong HLA-E binding affinity of the NKG2A receptor ectodomain with the activating signaling of the NKG2C receptor endodomain. We demonstrate that A/C Switch expression arms cytotoxic lymphocytes to overcome the inhibitory signaling of HLA-E and mediates cytotoxicity against liquid and solid cancer cell lines. The response was tuned by the receptor density at the cell surface and restricted to target cells expressing medium to high levels of HLA-E, thereby safeguarding against on-target, off-tumor toxicity. Moreover, A/C Switch transduced T cells were able to efficiently control tumor growth in an orthotopic glioblastoma *in vivo* model.

## Results and Discussion

### Design and expression of a novel switch receptor targeting HLA-E

We hypothesized that the inhibitory HLA-E checkpoint could be effectively overcome in T and NK cells through a novel switch receptor that combines the high HLA-E binding affinity of the extracellular domain of NKG2A with positive downstream signaling through the intracellular domain of NKG2C. Given that CD94 heterodimerization and the adaptor protein DAP12 are required for NKG2C signaling, we constructed a tetra-cistronic vector composed of our switch receptor, CD94, DAP12, and a truncated EGFR (EGFRt) reporter, hereafter referred to as the A/C Switch construct (Figures 1A-B).

**Figure 1:**
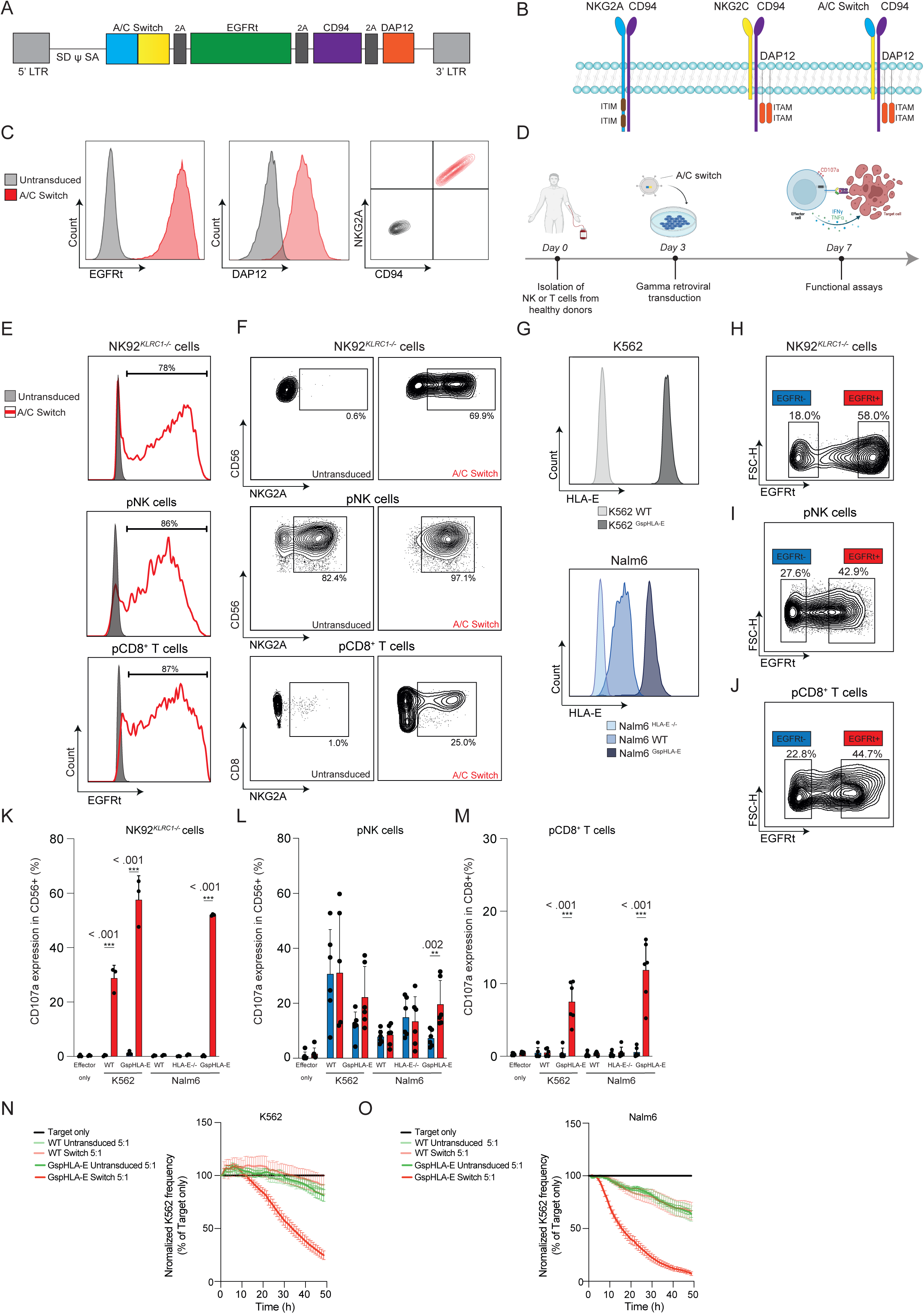
A/C Switch transduced effector cells respond to high, but not low levels of HLA-E on target cells. **A.** SFG-gammaretroviral tetra-cistronic vector map. **B.** Schematic depicting NKG2A (blue) and NKG2C (yellow) and A/C Switch (blue and yellow) receptors heterodimerized with CD94 (purple). **C.** Flow cytometry analysis 4 days post A/C Switch transduction of fibroblasts. Surface expression of EGFRt, NKG2A and CD94. Intracellular expression of DAP12. **D.** Schematic depicting experimental overview of functional in vitro assays. **E.** Transduction efficiency of A/C Switch in effector cells measured by EGFRt reporter (in red). **F.** NKG2A expression in untransduced and transduced NK92*^KLRC1-/-^*, pNK, and pCD8^+^ T cells. **G.** Variants of K562 cell line expressing either WT (light gray) or GspHLA-E (dark gray). Variants of Nalm6 cell line expressing either HLA-E^-/-^ (light blue), WT (blue) or GspHLA-E (dark blue). **H, I, J.** Gating on reporter EGFRt negative (blue) and EGFRt high (red) subsequently used for analysis in K, L and M. **K, L, M.** Degranulation assay comparing untransduced (blue) or A/C Switch transduced (red) NK92*^KLRC1-/-^*, pNK, and pCD8^+^ T cells against target cell lines from **G** as measured by surface CD107a expression. Effector cells were co-cultured with target cells at an E:T ratio of 5:1. Total of six donors in three independent experiments. *P* values were determined using Mann-Whitney test. **N, O.** Killing assay of A/C Switch transduced pCD8^+^ T cells against K562 and Nalm6 cell line variants labelled with nuclear red dye (NLR). Target only depicted in solid black line. WT target cell lines depicted in light green (against untransduced T cells) and light pink (against A/C Switch T cells). K562^GspHLA-E^ and Nalm6^GspHLA-E^ cells are shown in bold green (against untransduced T cells) and bold red (against A/C Switch T cells). Total of six donors in three independent experiments.

To determine whether all components of the A/C Switch construct were synthesized and expressed, we gamma-retrovirally transduced a fibroblast line that was negative for the four genes of interest. Flow cytometry staining identified transduced cells, which co-expressed NKG2A, CD94, DAP12, and EGFRt (Figure 1C), confirming the intended pattern of protein expression.

In agreement with previous studies, endogenous expression of NKG2A on human peripheral blood lymphocytes was variable and found in approximately 90% of CD56^bright^, 40% of CD56^dim^ NK cells, and 15% of CD56^+^ T cells. Conventional CD8^+^ T cells displayed on average 3% NKG2A expression, but in some individuals up to 20-30% (Supplementary Figure 1A), and it is well established that this checkpoint negatively regulates CD8^+^ T cell responses in the tumor microenvironment (Borst et al., 2020; Salome et al., 2022; van Montfoort et al., 2018). The NK92 cell line has a phenotype that resembles CD56^bright^ NK cells (Gunesch et al., 2019) with uniform NKG2A expression (Supplementary Figure 1B). Therefore, we used CRISPR/Cas9 editing to knock out *KLRC1*, which encodes NKG2A, in the NK92 cell line (NK92*^KLRC1-/-^*, Supplementary Figure 1B), enabling us to study the effects of A/C Switch in isolation from the inhibitory effects of endogenous NKG2A signaling (Supplementary Figure 1B). We transduced the A/C Switch receptor into NK92*^KLRC1-/-^* cells, primary human NK (pNK) cells, and primary human CD8^+^ (pCD8^+^ T) cells and performed functional assays on Day 7 (Figure 1D). Expression of the A/C Switch construct was detectable in NK92*^KLRC1-/-^*, pNK, and pCD8^+^ T cells using flow cytometry staining of the EGFRt reporter and NKG2A (Figures 1E-F). Expression of the critical recognition and signaling components of the A/C Switch were orthogonally verified in transduced pCD8^+^ T cells with western blot (Supplementary Figure 1C). A/C Switch expression did not affect the viability of NK92*^KLRC1-/-^*, pNK, or pCD8^+^ T cells compared to mock transduced cells (Supplementary Figure 1D).

### A/C Switch-transduced effector cells respond to high, but not low levels of HLA-E on target cells

We engineered target cell lines to investigate the specificity of the A/C Switch construct against a range of HLA-E expression levels (Figure 1G). Wild-type K562 (K562 WT) leukemia cells lack HLA-E expression, while wild-type Nalm6 (Nalm6 WT) leukemia cells express low levels of HLA-E. An HLA-E negative variant of Nalm6 was generated using CRISPR/Cas9 editing (Nalm6^HLA-E-/-^). To achieve high and stable expression of HLA-E, we introduced a single chain construct with the HLA-G leader sequence peptide (Gsp) linked to HLA-E (K562^GspHLA-E^ and Nalm6^GspHLA-E^). In A/C Switch-transduced NK92*^KLRC1-/-^* cells, NKG2A expression correlated with high levels of EGFRt, suggesting that those cells robustly express the A/C Switch receptor at the cell surface (Supplementary Figure 1E). We gated on cells with high levels of EGFRt to specifically evaluate the biology of the A/C Switch receptor in our functional assays (Figure 1H-J). The A/C Switch receptor triggered an increased degranulation response (CD107a) in A/C Switch-transduced NK92*^KLRC1-/-^* cells when stimulated with K562^GspHLA-E^ compared to HLA-E low K562 WT cells. The specificity of the response was further demonstrated against Nalm6^GspHLA-E^ cells, since there was no background activity against Nalm6 WT or Nalm6^HLA-E-/-^ cells (Figure 1K). Untransduced EGFRt negative NK92 cells remained unresponsive to both WT and HLA-E high targets.

Importantly, the A/C Switch restored degranulation and pro-inflammatory cytokine secretion of pNK cells against Nalm6^GspHLA-E^ cells at levels comparable to untransduced effectors challenged with Nalm6^HLA-E-/-^ cells, illustrating the switch function of the receptor (Figure 1L and Supplementary Figure 2A). In pNK cells, the A/C Switch competes for binding to HLA-E with high endogenous expression of inhibitory NKG2A receptors (Figure 1F). Indeed, depletion of NKG2A expressing pNK cells unleashed the full potential of the A/C Switch receptor against Nalm6^GspHLA-E^ cells (Supplementary Figure 2B).

In pCD8^+^ T cells, which express lower levels of competing endogenous NKG2A, the A/C Switch induced target-specific degranulation and synthesis of IFN-γ and TNF-α upon challenge with K562^GspHLA-E^ and Nalm6^GspHLA-E^, but not K562 WT, Nalm6 WT, or Nalm6^HLA-E-/-^ (Figure 1M and Supplementary Figure 2C). The same pattern was seen in long-term killing assays, where A/C Switch-transduced pCD8^+^ T cells specifically eliminated K562^GspHLA-E^ and Nalm6^GspHLA-E^ (Figures 1N-O). The distinct specificity and threshold response observed in pCD8^+^ T cells motivated our subsequent focus on A/C Switch-transduced T cells as a platform for extending the concept to solid tumors and proof-of-principle experiments in mice.

### A/C Switch-transduced T cells mediate cytotoxicity against liquid and solid tumor models with induced endogenous HLA-E expression

We next turned our focus to the U251 glioblastoma cell line, an established solid tumor model to investigate the efficacy of chimeric antigen receptor (CAR) engineered T cells in glioblastoma (Choi et al., 2019; Larson et al., 2022). In culture, U251 expressed low levels of endogenous HLA-E, which could be induced to medium levels with IFN-γ treatment (Supplementary Figure 3A). To model high levels of expression, we made a U251^GspHLA-E^ variant for *in vitro* and *in vivo* studies. A/C Switch engineered pCD8^+^ T cells showed some degree of recognition of U251 WT cells, but killing was significantly increased against targets expressing high levels of HLA-E (Figure 2A), in line with the data obtained in the leukemia model. In a competitive killing assay, where equal numbers of U251 WT and U251^GspHLA-E^ cells were labelled with different concentrations of cell trace violet (CTV), combined, and incubated with A/C Switch-transduced pCD8^+^ T cells or untransduced controls, U251^GspHLA-E^ were preferentially eliminated in a target-specific manner (Figure 2B and Supplementary Figure 3B).

**Figure 2:**
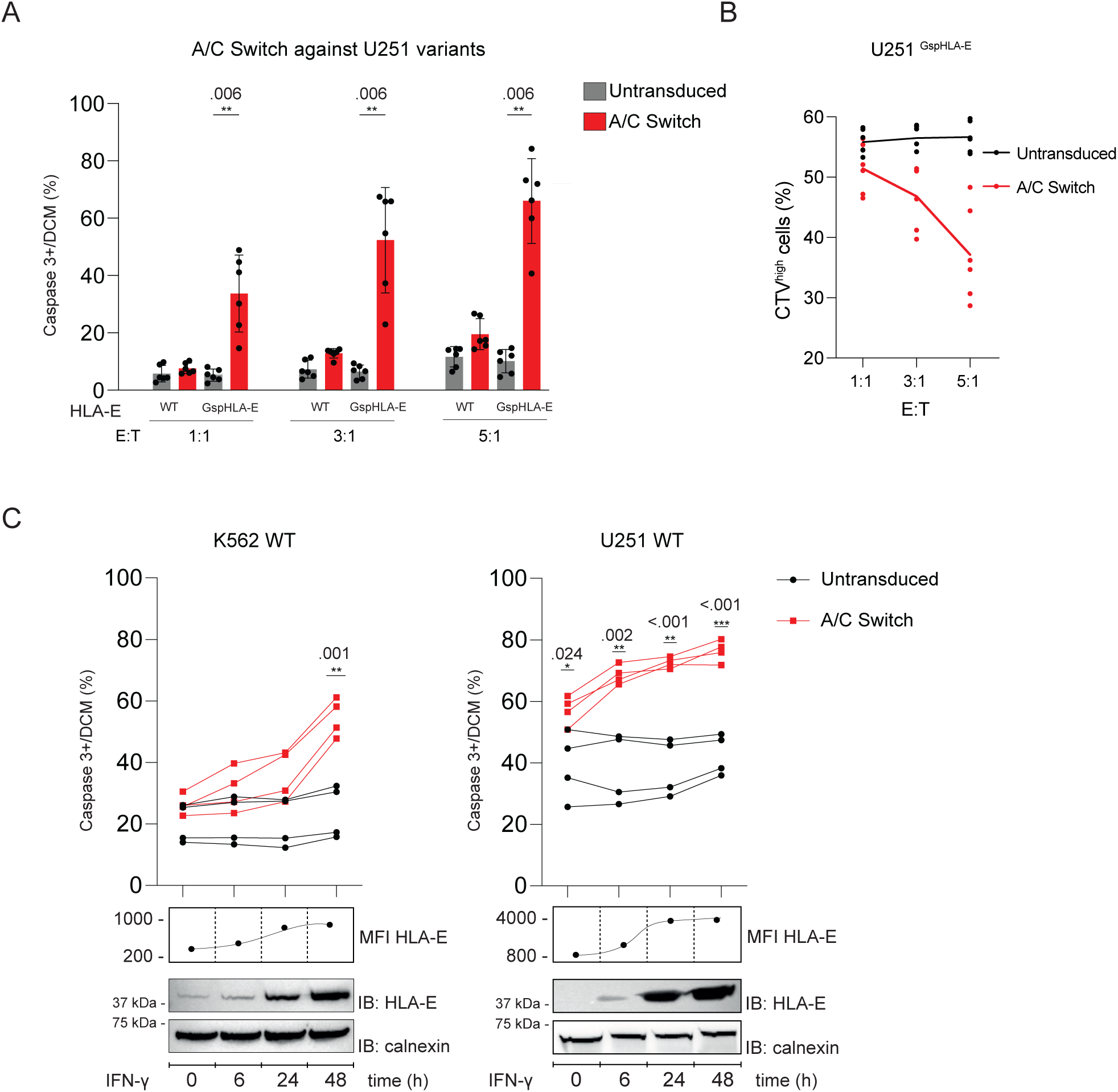
A/C Switch transduced pCD8^+^ T cells mediate cytotoxicity against liquid and solid tumor models with induced endogenous HLA-E expression. **A.** Cytotoxicity assay of A/C Switch transduced pCD8^+^ T cells (red) compared to untransduced (gray) at three different E:T ratios co-cultured with U251 glioblastoma cell line variants. Cell death measured as caspase 3^+^ /Dead Cell Marker percent positive cells (%). Total of six donors in two independent experiments. **B.** Killing assay of U251^GspHLA-E^ target cells (CTV^high^ labeled) by A/C Switch T cells (red) and untransduced (black) at different E:T ratios. Total of six donors. **C.** Killing assay of K562 WT and U251 WT cells displaying IFN-γ-mediated HLA-E upregulation by A/C Switch pCD8^+^ T cells (red) and untransduced (black) at 5:1 E:T ratio. Target cells were treated with 20ng/ml IFN-γ for 48hr. Endogenous HLA-E expression is shown by western blot and flow cytometry analyses (MFI). Total of four donors in two independent experiments. *P* values were determined using Mann-Whitney test.

Thus far, we found that the A/C Switch induced potent functional responses to high levels of HLA-E expression but remained unresponsive to low levels. This is an important prerequisite, since HLA-E is ubiquitously expressed by normal cells, albeit mostly at low levels. Indeed, A/C Switch T cells did not induce on-target off-tumor killing of normal cell types derived from cell lineages reported to have the highest endogenous HLA-E expression in the Human Protein Atlas, including monocytes, adipocytes, NK cells, endothelial cells, T cells, and B cells, even when treated with IFN-γ (Supplementary Figure 3C-D). These results suggest that the A/C Switch construct can be used to target elevated cancer-associated HLA-E, while avoiding on-target off-tumor effects in normal tissues, a major challenge in adoptive T cell therapies (Flugel et al., 2023).

Notably, treatment of K562 cells and U251 cells with 20 ng/ml of IFN-γ, in the range of serum concentrations observed after CAR T cell activation (Jain et al., 2023), increased the HLA-E expression and sensitized both lines to A/C Switch-transduced T cells (Figure 2C, Supplementary Figure 3E). The ability of A/C Switch pCD8^+^ T cells to kill tumor cells exposed to IFN-γ is relevant in the context of induced resistance to CAR T cell therapy (Hamieh et al., 2019; Hamieh et al., 2023; Majzner and Mackall, 2018). In contrast to conventional CAR-T cells, A/C Switch-bearing T cells are poised to generate a response-amplifying positive feedback loop through A/C Switch-triggered production of IFN-γ and the induction of the A/C Switch ligand, HLA-E (Barrett et al., 2004; Gustafson and Ginder, 1996; Malmberg et al., 2002; Nguyen et al., 2009).

### *In vivo* cytotoxicity is modulated by target cell HLA-E and T cell A/C Switch levels

To test the *in vivo* efficacy of A/C Switch-transduced primary human T cells, we used an orthotopic glioblastoma mouse model (Figure 3A). In the absence of a CAR or functional TCR (Supplementary Figure 1F), A/C Switch-transduced T cells were able to fully eradicate U251^GspHLA-E^ tumors, enabling survival past 100 days (Figures 3B-D). To determine whether the cytotoxic function of the A/C Switch receptor could be modulated, we reduced transduction efficiency to ≤30%, which in turn reduced A/C Switch construct expression density (about 2.5-fold relative MFI reduction) (Figure 3E). A/C Switch T cells with lower surface density of the receptor exhibited decreased ability to control orthotopic glioblastomas (Figures 3F-G, median survival 34 vs 43 days, p<0.002), consistent with decreases in effector cytokine secretion observed *in vitro* with lower A/C Switch per T cell (Supplementary Figure 3F-G). In line with the *in vitro* data, T cells expressing high A/C Switch levels were unable to control HLA-E low U251 WT orthotopic tumors (Figure 3H).

**Figure 3:**
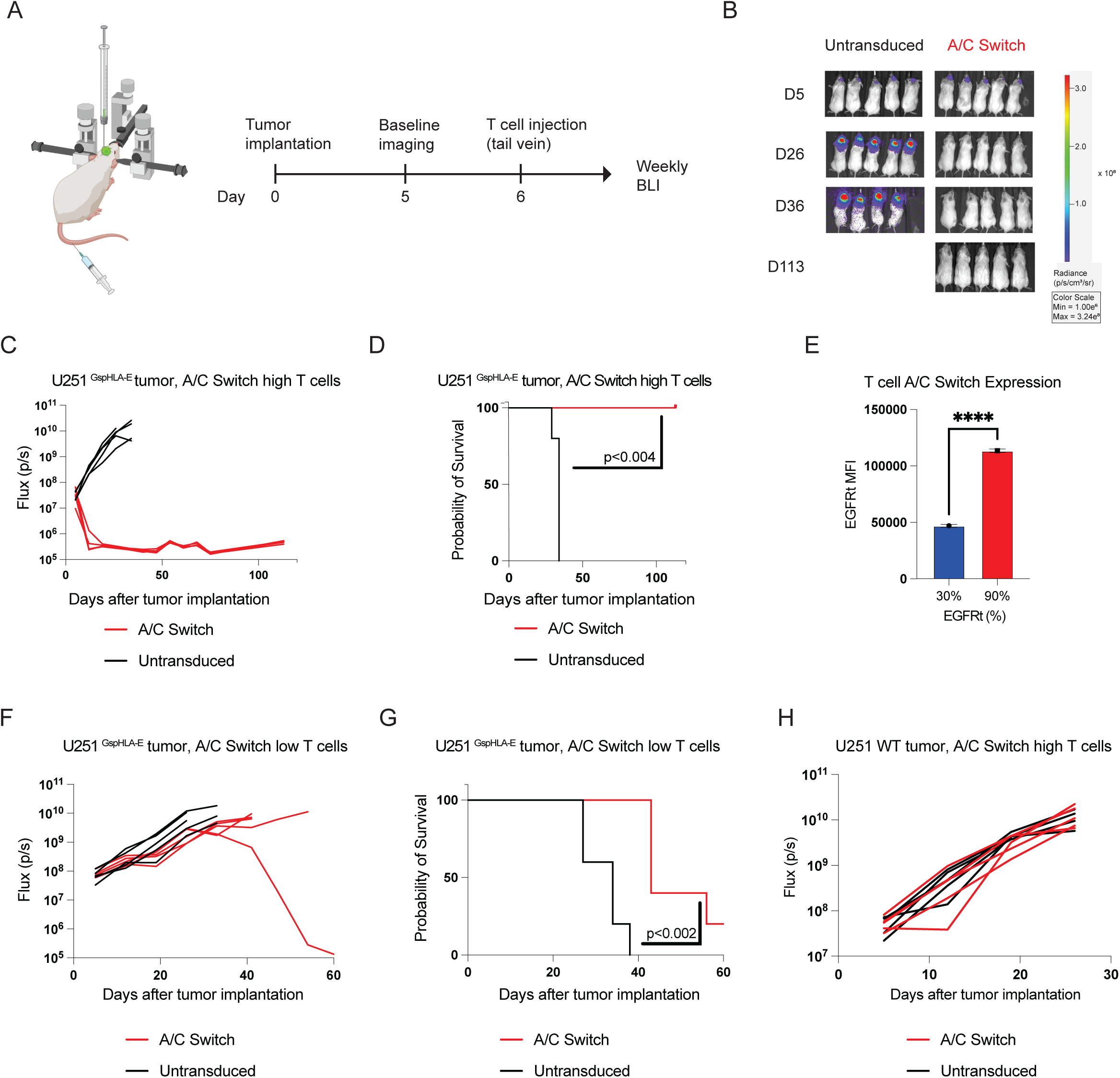
*In vivo* cytotoxicity is modulated by target cell HLA-E and T cell A/C Switch levels. **A.** Schematic of experimental setup for results shown in B-H. **B.** Bioluminescent imaging of NSG mice bearing U251^GspHLA-E^ ffLuc+ orthotopic glioblastomas treated with A/C Switch high (red) or untransduced (black) T cells. **C and D.** Tumor burden represented by bioluminescence and survival analysis of NSG mice bearing U251^GspHLA-E^ glioblastomas treated with A/C Switch (red) or untransduced T cells (black). **E.** Cell surface expression of EGFRt A/C Switch T cells, low 30% (blue) and high 90% (red), p<0.0001 by unpaired two-tailed T test. **F and G.** Tumor burden represented by bioluminescence and survival analysis of NSG mice bearing U251^GspHLA-E^ treated with A/C Switch low (red) or untransduced (black) T cells. **H.** Tumor burden represented by bioluminescence of NSG mice bearing U251 WT treated with A/C Switch high (red) or untransduced (black) T cells. Survival was analyzed using Kaplan-Meier methodology and statistical comparison of survival curves was performed using the Log-rank test.

These results indicate that the current A/C Switch construct has a relatively broad safety margin, providing a therapeutic window to minimize toxicity against normal cells with low HLA-E expression, while efficiently targeting tumor cells with medium to high HLA-E, whether spontaneously or induced by proinflammatory cytokines such as IFN-γ in the tumor microenvironment. In its current version, the A/C Switch receptor drives T cells to mediate robust cytotoxicity when both the A/C Switch and HLA-E are present in high abundance. This may be related to the strength of DAP12 signaling in T cells. Future studies will determine whether this feature may be tunable through engineering approaches that use alternative or calibrated signaling proteins or augment the current construct’s sensitivity through combinatorial strategies using CARs.

In conclusion, the A/C Switch described herein represents a cell intrinsic approach to relieve the immune inhibition of the HLA-E/NKG2A checkpoint that may be used alone or in combination with other targeting modalities.

## Materials and Methods

### Reagents used in the study

Reagents used are listed in Supplementary Table 1.

### Cells

#### Primary cells

For *in vivo* studies buffy coats from anonymous healthy donors were purchased from the New York Blood Center (institutional review board-exempted) and for *in vitro* studies peripheral blood samples from healthy donors were obtained from Oslo University Hospital as approved by the regional ethical committee (REK 2018/2485 and 2018/2482. PBMCs were separated by centrifugation using a density gradient medium (Lymphoprep, Serumwerk). Different subsets, including NK cells, pan T cells, CD8^+^ or CD4^+^ T cells, CD14^+^ and CD19^+^ cells were isolated from PBMCs using the corresponding kits (Miltenyi) and according to manufacturer’s protocol. Cells were maintained in RPMI media (Gibco, ThermoFisher) supplemented with 10% FCS (Sigma-Aldrich) and PenStrep (Sigma-Aldrich) at 37°C, 5% CO_2_.

#### Cell lines

NK92 cells were purchased (ATCC) and maintained in MEMα, nucleosides (ThermoFisher) supplemented with 2mM L-glut, 12.5% horse serum (ThermoFisher), 12.5% FCS (Sigma-Aldrich), 1% PenStrep and 100IU/ml interleukin-2 (ThermoFisher). K562 cells, a chronic myeloid leukemia cell line, and Nalm6 cells, a B cell acute lymphoblastic leukemia cell line, were purchased from ATCC. K562 variants and Nalm6 variants were maintained in RPMI media (ThermoFisher) supplemented with 10% FCS (Sigma-Aldrich) and PenStrep (Sigma-Aldrich) at 37°C, 5% CO_2_. U251 glioblastoma cells were purchased from Millipore Sigma and maintained in DMEM supplemented with 10% FCS (Sigma-Aldrich) and PenStrep (Sigma-Aldrich) at 37°C, 5% CO_2_. NK92*^KLRC1-/-^* cells and Nalm6^HLA-E-/-^ cells were generated using CRISPR/Cas9 editing. Wild-type K562 and U251 as well as Nalm6^HLA-E-/-^ cells were used to generate K562^GspHLA-E^, U251^GspHLA-E^, and Nalm6^GspHLA-E^ by overexpressing HLA-E*01:01 as a single chain construct covalently linked to β2m and to the HLA-G leader sequence peptide VMAPRTLFL (Gsp), as described previously (https://www.biorxiv.org/content/10.1101/2023.10.09.557143v1).

Adipocytes were differentiated from human adipose-derived stem cells (ADSCs, ThermoFisher), which were obtained from human lipoaspirate tissue and cryopreserved from primary culture. ADSCs/adipocyctes were maintained in MesenPRO RS™ Basal Medium supplemented with MesenPRO RS™ Growth Supplement according to suppliers’ protocol at 37°C, 5% CO_2_. Human Umbilical Vein Endothelial Cells (HUVEC) were obtained from ThermoFisher and maintained in Human Large Vessel Endothelial Cell Basal Medium (ThermoFisher) according to manufacturer’s protocol.

### Gammaretroviral vector construct and production

Plasmids encoding the SFG-γ retroviral vector (Riviere et al., 1995) were prepared using standard molecular biology techniques as described previously (Brentjens et al., 2003; Maher et al., 2002). Our tetracistronic construct was cloned into the SFG-γ-retroviral vector as outlined in Figure 1A. A/C Switch was co-expressed with a truncated EGFR (EGFRt) reporter, CD94, and DAP12. VSV-G pseudotyped retroviral supernatants derived from transduced gpg29 fibroblasts (H29) were used to generate stable retroviral-producing cell lines (Gong et al., 1999).

### Gammaretroviral transduction

#### Activation of effector cells prior to transduction

NK cells were activated by co-culturing NK cells with irradiated K562 cells expressing membrane bound interleukin-21 and 4-1BBL at 1:1 ratio in RPMI (Gibco, Thermo Fisher) supplemented with 10% FCS (Sigma-Aldrich), Pen-Strep (Sigma-Aldrich) and 200UI/mL of interleukin-2 (ThermoFisher) for 3 days. T cells were activated with Human T-Activator CD3/CD28 Dynabeads (Thermo Fisher) at a 1:1 bead:cell ratio in RPMI (Gibco, Thermo Fisher) supplemented with 10% FCS (Sigma-Aldrich), PenStrep (Sigma-Aldrich), and 5ng/ml of human recombinant interleukin-7 (Peprotech) and 5ng/ml human recombinant interleukin-15 (Miltenyi) for 3 days.

#### Transduction

Effector cells (NK92, T cells, and NK cells) were transduced with retroviral supernatants by centrifugation on RetroNectin (Takara) coated plates. Transduction efficiencies were determined 3 days later by flow cytometry and used in *in-vitro* or *in-vivo* experiments as outlined in Figure 1D.

### gRNA, Cas9 protein and RNP formation

#### T cells

The TCRα subunit constant gene (*TRAC)* was targeted with sgRNA (target sequence: 5’ - CAGGGTTCTGGATATCTGT) (Eyquem et al., 2017). *TRAC* sgRNA was purchased from Synthego with 20 -O-methyl 30 -phosphorothioate modifications in the first and last three nucleotides. Guide RNA was resuspended with TE buffer to a concentration of 40mM. Cas9 protein (40mM) was obtained from QB3-Berkeley Macrolab core facility. RNP formation was prepared by mixing Cas9 protein and *TRAC* gRNA at 1:1 molar ratio, incubating at 37°C for 15 minutes. CRISPR/Cas9 editing was used to knockout TRAC in T cells for *in vivo* experiments. 48h post T cell activation, the CD3/CD28 beads were magnetically removed, and T cells were electroporated and transfected with TRAC RNP using a 4D-Nucleofector device (Lonza) and P3 Primary Cell Solution according to manufacturer protocol (Lonza). Following electroporation, cells were incubated in culture medium at 1×10^6^ cells/ml. Twenty-four hours post electroporation, T cells were γ-retrovirally transduced as described previously. TRAC edited and SFG γ-retrovirally-transduced T cells were maintained in RPMI (ThermoFisher) supplemented with 10% FCS (Sigma-Aldrich), PenStrep (Sigma-Aldrich), and 5ng/ml human recombinant interleukin-7 (Peprotech) and 5ng/ml human recombinant interleukin-15 (Miltenyi) at 1×10^6^ −1.5×10^6^ cells/ml. *Nalm6 and NK92:*

CRISPR/Cas9 editing was used to disrupt *HLA-E* in Nalm6 and *KLRC1* in NK92 cells. Nalm6 and NK92 cells were resuspended in SF buffer (Lonza) and mixed with HLA-E RNP or NKG2A RNP in a total volume of 100ul. Following electroporation, Nalm6 and NK92 cells were incubated and expanded in their respective medium as described above.

HLA-E gRNA target sequence:

5’-ACAACGACGCCGCGAGTCCG

NKG2A gRNAs target sequences:

5’-AAGCTTCTCAGGATTTTCAA, 5’-AGGCAGCAACGAAAACCTAA,

5’-ACTGCAGAGATGGATAACCA)

### Flow cytometry assays

Cells were stained following standard flow cytometry procedures. Briefly, cells were collected in a V-bottom 96-well plate (Falcon) and resuspended in 45 μl of FACS buffer (0.5% BSA, 10% FBS in PBS) containing antibodies for extracellular markers. Cells were stained for 30 minutes at room temperature in the dark. For assays that required fixation, cells were fixed using Cytofix/Cytoperm (BD) following manufacturer’s protocol. Full list of antibodies is available in Table 1.

To determine HLA-E expression in target cell lines, cells were incubated with anti-HLA-E antibody and a Live/Dead Aqua dye (ThermoFisher) for 30 minutes at 4°C. Samples were then fixed and acquired on flow cytometer.

#### Functional assays

Functional assays were performed at 37°C in RPMI (ThermoFisher) supplemented with 10% FCS (Sigma-Aldrich) and PenStrep (Sigma-Aldrich). Effector cells were incubated with K562 and Nalm6 variants for 4 hours at a ratio of 5:1 with addition of Brefeldin A (GolgiPlug, BD Biosciences). Next, cells were centrifuged at 300 x *g*, surface stained with lineage markers and anti-CD107a antibodies for assessing degranulation. Cells were then fixed with Cytofix/Cytoperm (BD Biosciences) and stained for intracellular cytokines (IFN-γ, TNF-α).

#### Competitive killing assay

U251 cell variants (U251 WT and U251 ^GspHLA-E^) were labeled with two different CellTrace Violet (ThermoFisher) concentrations (0.1 μM, 5.0 μM) and seeded in a flat-bottom 96-well plate (ThermoFisher). After 4 hours, transduced effector cells were added at different E:T ratios and RED-DEVD-FMK (Abcam) reagent was added to detect active caspase-3. Effector and targets were co-cultured for 18 hours at 37°C, 5% CO_2_. Post incubation, cells were surface stained with fluorochrome-conjugated antibodies and Fixable Viability Stain 780 (ThermoFisher).

#### Normal cell killing assay

CD3^+^, CD56^+^, CD14^+^, and CD19^+^ cells were isolated from PBMCs as described above. HUVEC and adipocyte cells were purchased from ThermoFisher. Cells were transferred to 96-well plates and treated with 20ng/ml of IFN-γ (R&D Systems) for 48 hours prior to co-culture with T cells. RED-DEVD-FMK (Abcam) reagent and effector T cells (untransduced and A/C Switch) were added at 5:1 E:T ratio and co-cultured with targets for 18 hours at 37°C, 5% CO_2._ Post incubation, cells were surface stained with fluorochrome-conjugated antibodies and Fixable Viability Stain 780 (ThermoFisher).

All flow cytometry samples were acquired using the BD Symphony instrument (BD Biosciences) and analyzed using FlowJo v.10.2 software (FlowJo).

### Western blotting

For assessment of NKG2A, DAP12 and HLA-E protein expression, one million cells per condition were lysed in RIPA buffer (ThermoFisher) supplemented with 1x HALT protease & phosphatase inhibitor cocktail (ThermoFisher) for 30 minutes on ice, followed by centrifugation. Supernatants were mixed with 4x NuPage loading buffer (ThermoFisher) and 10x Sample Reducing Agent (ThermoFisher), and run on a 4-12% Bis-Tris gel (ThermoFisher) then transferred to a iBlot2 PVDF membrane (ThermoFisher). Secondaries antibodies included goat anti-mouse (Cell Signaling Tech.) and goat anti-rabbit (Cell Signaling Tech.). Blocking buffer and antibodies were diluted in 5% non-fat dry milk (Sigma-Aldrich) in TBS-Tween 0.1%. Membranes were developed using SuperSignal substrate (ThermoFisher) in iBright (ThermoFisher). For increasing HLA-E expression levels, K562 and Nalm6 WT cells were incubated with 20 ng/mL IFN-γ up to 48 hours.

### IncuCyte killing assay

Specific tumor lysis was measured in real-time using the IncuCyte S3 platform as described previously (Haroun-Izquierdo et al., 2022). Briefly, target cells stably expressing NucLight Red (Essen Biosciences) were plated and rested overnight and subsequently co-cultured with A/C Switch-transduced effector cells at different E:T ratios. Images (3/well) from at least three technical replicates for each condition were acquired every 90 min for 48 hours, using a ×10 objective lens and analyzed by IncuCyte Controller v2020A (Essen Biosciences). Graphed readouts represent percentage live target cells (based on NLR expression).

#### NKG2A depletion killing assay

PBMCs from healthy donors were screened and donors were selected based on a minimum of 50% NKG2A positive expression in the CD3^-^CD56^+^ compartment. Bulk NK cells were isolated as described previously from PBMCs. To deplete NKG2A from freshly isolated bulk NK cells, a biotinylated anti-NKG2A antibody (Miltenyi Biotec) and Anti-Biotin Microbeads (Miltenyi Biotec) were used for labeling and depletion of NKG2A^+^ cells. Transduction with A/C Switch was performed as previously described and detailed in Figure 1D and used in IncuCyte assays as described above.

### Mouse orthotopic glioblastoma model

All animal experiments were approved by the Memorial Sloan Kettering Cancer Center (MSK) Institutional Animal Care and Use Committee (IACUC). NSG Mice (Jackson Labs, USA) were anesthetized with ketamine and dexmedetomidine. Using sterile technique, the cranium was accessed to drill a burr hole 2 mm lateral to bregma and 5 x 104 U251 cells bearing firefly luciferase GFP were implanted at a depth of 3 mm. Perioperative anesthesia was provided with meloxicam. Five days after tumor implantation, bioluminescent imaging (BLI) using the IVIS Spectrum in vivo imaging system (PerkinElmer, USA) was initiated on a weekly basis. Six days after tumor implantation, untransduced or A/C Switch T cells were injected via tail vein. Untransduced T cells were always infused at a dose equivalent to the total number of T cells in A/C Switch conditions (untransduced + A/C Switch). Mice were followed daily for signs of clinical deterioration, such as lethargy and skin pallor, requiring euthanasia.

### Statistical analyses

Details on statistics can be found in figure legends. All statistical analyses were performed using GraphPad Prism software version 10.0.1 (GraphPad). *P* values *<*0.05 were considered statistically significant. The Kaplan Meier method was used for survival representations and log-rank test was used to compare survival differences between the groups. The statistical test used for each figure is described in the corresponding figure legend.

## Supplementary Figure Legends

**Figure S1: NKG2A expression and A/C switch transduction.**

**A.** Expression of NKG2A in a cohort of 202 donors in (left) CD56^bright^ and CD56^dim^ NK cells, (middle) CD56^+^ T cells, and (right) conventional CD3^+^CD56^-^, CD3^+^CD56^-^CD8^+^ and CD3^+^CD56^-^ CD8^-^CD57^-^ T cells. **B.** Flow cytometry analysis of NKG2A and CD56 expression in NK92 WT (orange) and NK92 *^KLRC1-/-^* cells (blue). **C.** Western blot of each gene from the tetra-cistronic construct present in A/C Switch transduced pCD8^+^ T cells. Calnexin was used as loading control. **D.** Flow cytometry viability analysis of NK92*^KLRC1-/-^*, pNK or pCD8^+^ T cells when transduced with A/C Switch as shown by expression of dead cell marker. **E.** Co-expression shown as bivariate plot of NKG2A and EGFRt in A/C Switch transduced NK92*^KLRC1-/-^* cells. **F.** *TRAC* knockout efficiency measured by flow cytometry analysis of CD3 surface expression on T cells. Total of four donors in two independent experiments.

**Figure S2: Functional response of A/C Switch transduced primary NK and pCD8 T cells**

**A and C.** Intracellular levels of TNF-α and IFN-γ in A/C Switch transduced pNK and pCD8^+^ T EGFRt high (red) and untransduced (blue) cells when co-cultured with target cells (K562 WT, K562^GspHLA-E^, Nalm6, Nalm6^HLA-E-/-^, and Nalm6^GspHLA-E^). Total of six donors in three independent experiments. *P* values were determined using Mann-Whitney test. **B.** Depletion of NKG2A^+^ cells from bulk pNK cells. Left bivariate plot shows CD56/NKG2A expression pre-sort and bivariate plot to the right shows CD56/NKG2A expression post-sort. NKG2A depleted (red) and bulk (gray) A/C Switch pNK cells used in killing assay against Nalm6^GspHLA-E^. Total of three donors in two independent experiments.

**Figure S3: Safety profile and therapeutic window of A/C Switch transduced pCD8 T cells**

**A.** Histogram of HLA-E expression in U251 WT (red), U251 WT^IFN-γ^ (blue) and U251^GspHLA-E^ (orange) cells. **B.** Competitive killing assay with two target cell lines, U251 WT (CTV low) and U251^GspHLA-E^ (CTV high). Untransduced (top row) or A/C Switch transduced T cells (bottom row) were co-cultured with U251 WT and U251^GspHLA-E^ at three different effector-to-target cell ratios, 1:1, 3:1, and 5:1. **C.** HLA-E single cell RNA number of transcripts per million (nTPM) from Human Protein Atlas. **D.** A/C Switch (red) or untransduced (gray) T cells were co-cultured with the top five highest HLA-E expressing cell subsets from **C**, with (slashed pattern) and without (solid) IFN-γ treatment (20ng/ml) for 48h. **E.** U251 WT cells treated using the same conditions as in D; data extracted from experiment performed in Figure 2C. **F and G.** Expression of cytotoxic markers (surface CD107a and intracellular IFN-γ and TNF-α in untransduced (gray), A/C Switch low (40%), and A/C Switch high (90%) T cells co-cultured with Nalm6^GspHLA-E^ and K562^GspHLA-E^. *P* values were determined using Mann-Whitney test.

## References

Andersson, E., I. Poschke, L. Villabona, J.W. Carlson, A. Lundqvist, R. Kiessling, B. Seliger, and G.V. Masucci. 2016. Non-classical HLA-class I expression in serous ovarian carcinoma: Correlation with the HLA-genotype, tumor infiltrating immune cells and prognosis. Oncoimmunology 5:e1052213.

Andre, P., C. Denis, C. Soulas, C. Bourbon-Caillet, J. Lopez, T. Arnoux, M. Blery, C. Bonnafous, L. Gauthier, A. Morel, B. Rossi, R. Remark, V. Breso, E. Bonnet, G. Habif, S. Guia, A.I. Lalanne, C. Hoffmann, O. Lantz, J. Fayette, A. Boyer-Chammard, R. Zerbib, P. Dodion, H. Ghadially, M. Jure-Kunkel, Y. Morel, R. Herbst, E. Narni-Mancinelli, R.B. Cohen, and E. Vivier. 2018. Anti-NKG2A mAb Is a Checkpoint Inhibitor that Promotes Anti-tumor Immunity by Unleashing Both T and NK Cells. Cell 175:1731–1743 e1713.

Barrett, D.M., K.S. Gustafson, J. Wang, S.Z. Wang, and G.D. Ginder. 2004. A GATA factor mediates cell type-restricted induction of HLA-E gene transcription by gamma interferon. Mol Cell Biol 24:6194–6204.

Borst, L., M. Sluijter, G. Sturm, P. Charoentong, S.J. Santegoets, M. van Gulijk, M.J. van Elsas, C. Groeneveldt, N. van Montfoort, F. Finotello, Z. Trajanoski, S.M. Kielbasa, S.H. van der Burg, and T. van Hall. 2022. NKG2A is a late immune checkpoint on CD8 T cells and marks repeated stimulation and cell division. Int J Cancer 150:688–704.

Borst, L., S.H. van der Burg, and T. van Hall. 2020. The NKG2A-HLA-E Axis as a Novel Checkpoint in the Tumor Microenvironment. Clin Cancer Res 26:5549–5556.

Braud, V.M., H. Aldemir, B. Breart, and W.G. Ferlin. 2003. Expression of CD94-NKG2A inhibitory receptor is restricted to a subset of CD8+ T cells. Trends Immunol 24:162–164.

Braud, V.M., D.S. Allan, C.A. O’Callaghan, K. Soderstrom, A. D’Andrea, G.S. Ogg, S. Lazetic, N.T. Young, J.I. Bell, J.H. Phillips, L.L. Lanier, and A.J. McMichael. 1998. HLA-E binds to natural killer cell receptors CD94/NKG2A, B and C. Nature 391:795–799.

Brentjens, R.J., J.B. Latouche, E. Santos, F. Marti, M.C. Gong, C. Lyddane, P.D. King, S. Larson, M. Weiss, I. Riviere, and M. Sadelain. 2003. Eradication of systemic B-cell tumors by genetically targeted human T lymphocytes co-stimulated by CD80 and interleukin-15. Nat Med 9:279–286.

Cappell, K.M., and J.N. Kochenderfer. 2023. Long-term outcomes following CAR T cell therapy: what we know so far. Nat Rev Clin Oncol 20:359–371.

Carretero, M., C. Cantoni, T. Bellon, C. Bottino, R. Biassoni, A. Rodriguez, J.J. Perez-Villar, L. Moretta, A. Moretta, and M. Lopez-Botet. 1997. The CD94 and NKG2-A C-type lectins covalently assemble to form a natural killer cell inhibitory receptor for HLA class I molecules. Eur J Immunol 27:563–567.

Carretero, M., G. Palmieri, M. Llano, V. Tullio, A. Santoni, D.E. Geraghty, and M. Lopez-Botet. 1998. Specific engagement of the CD94/NKG2-A killer inhibitory receptor by the HLA-E class Ib molecule induces SHP-1 phosphatase recruitment to tyrosine-phosphorylated NKG2-A: evidence for receptor function in heterologous transfectants. Eur J Immunol 28:1280–1291.

Choi, B.D., X. Yu, A.P. Castano, A.A. Bouffard, A. Schmidts, R.C. Larson, S.R. Bailey, A.C. Boroughs, M.J. Frigault, M.B. Leick, I. Scarfo, C.L. Cetrulo, S. Demehri, B.V. Nahed, D.P. Cahill, H. Wakimoto, W.T. Curry, B.S. Carter, and M.V. Maus. 2019. CAR-T cells secreting BiTEs circumvent antigen escape without detectable toxicity. Nat Biotechnol 37:1049–1058.

Dubrot, J., P.P. Du, S.K. Lane-Reticker, E.A. Kessler, A.J. Muscato, A. Mehta, S.S. Freeman, P.M. Allen, K.E. Olander, K.M. Ockerman, C.H. Wolfe, F. Wiesmann, N.H. Knudsen, H.W. Tsao, A. Iracheta-Vellve, E.M. Schneider, A.N. Rivera-Rosario, I.C. Kohnle, H.W. Pope, A. Ayer, G. Mishra, M.D. Zimmer, S.Y. Kim, A. Mahapatra, H. Ebrahimi-Nik, D.T. Frederick, G.M. Boland, W.N. Haining, D.E. Root, J.G. Doench, N. Hacohen, K.B. Yates, and R.T. Manguso. 2022. In vivo CRISPR screens reveal the landscape of immune evasion pathways across cancer. Nat Immunol 23:1495–1506.

Eyquem, J., J. Mansilla-Soto, T. Giavridis, S.J. van der Stegen, M. Hamieh, K.M. Cunanan, A. Odak, M. Gonen, and M. Sadelain. 2017. Targeting a CAR to the TRAC locus with CRISPR/Cas9 enhances tumour rejection. Nature 543:113–117.

Flugel, C.L., R.G. Majzner, G. Krenciute, G. Dotti, S.R. Riddell, D.L. Wagner, and M. Abou-El-Enein. 2023. Overcoming on-target, off-tumour toxicity of CAR T cell therapy for solid tumours. Nat Rev Clin Oncol 20:49–62.

Gong, M.C., J.B. Latouche, A. Krause, W.D. Heston, N.H. Bander, and M. Sadelain. 1999. Cancer patient T cells genetically targeted to prostate-specific membrane antigen specifically lyse prostate cancer cells and release cytokines in response to prostate-specific membrane antigen. Neoplasia 1:123–127.

Gooden, M., M. Lampen, E.S. Jordanova, N. Leffers, J.B. Trimbos, S.H. van der Burg, H. Nijman, and T. van Hall. 2011. HLA-E expression by gynecological cancers restrains tumor-infiltrating CD8(+) T lymphocytes. Proc Natl Acad Sci U S A 108:10656–10661.

Gunesch, J.T., L.S. Angelo, S. Mahapatra, R.P. Deering, J.E. Kowalko, P. Sleiman, J.W. Tobias, L. Monaco-Shawver, J.S. Orange, and E.M. Mace. 2019. Genome-wide analyses and functional profiling of human NK cell lines. Mol Immunol 115:64–75.

Gustafson, K.S., and G.D. Ginder. 1996. Interferon-gamma induction of the human leukocyte antigen-E gene is mediated through binding of a complex containing STAT1alpha to a distinct interferon-gamma-responsive element. J Biol Chem 271:20035–20046.

Hamieh, M., A. Dobrin, A. Cabriolu, S.J.C. van der Stegen, T. Giavridis, J. Mansilla-Soto, J. Eyquem, Z. Zhao, B.M. Whitlock, M.M. Miele, Z. Li, K.M. Cunanan, M. Huse, R.C. Hendrickson, X. Wang, I. Riviere, and M. Sadelain. 2019. CAR T cell trogocytosis and cooperative killing regulate tumour antigen escape. Nature 568:112–116.

Hamieh, M., J. Mansilla-Soto, I. Riviere, and M. Sadelain. 2023. Programming CAR T Cell Tumor Recognition: Tuned Antigen Sensing and Logic Gating. Cancer Discov 13:829–843.

Haroun-Izquierdo, A., M. Vincenti, H. Netskar, H. van Ooijen, B. Zhang, L. Bendzick, M. Kanaya, P. Momayyezi, S. Li, M.T. Wiiger, H.J. Hoel, S.Z. Krokeide, V. Kremer, G. Tjonnfjord, S. Berggren, K. Wikstrom, P. Blomberg, E. Alici, M. Felices, B. Onfelt, P. Hoglund, B. Valamehr, H.G. Ljunggren, A. Bjorklund, Q. Hammer, L. Kveberg, F. Cichocki, J.S. Miller, K.J. Malmberg, and E. Sohlberg. 2022. Adaptive single-KIR(+)NKG2C(+) NK cells expanded from select superdonors show potent missing-self reactivity and efficiently control HLA-mismatched acute myeloid leukemia. J Immunother Cancer 10:

Herbst, R.S., M. Majem, F. Barlesi, E. Carcereny, Q. Chu, I. Monnet, A. Sanchez-Hernandez, S. Dakhil, D.R. Camidge, L. Winzer, Y. Soo-Hoo, Z.A. Cooper, R. Kumar, J. Bothos, C. Aggarwal, and A. Martinez-Marti. 2022. COAST: An Open-Label, Phase II, Multidrug Platform Study of Durvalumab Alone or in Combination With Oleclumab or Monalizumab in Patients With Unresectable, Stage III Non-Small-Cell Lung Cancer. J Clin Oncol 40:3383–3393.

Houchins, J.P., L.L. Lanier, E.C. Niemi, J.H. Phillips, and J.C. Ryan. 1997. Natural killer cell cytolytic activity is inhibited by NKG2-A and activated by NKG2-C. J Immunol 158:3603–3609.

Hrbac, T., A. Kopkova, F. Siegl, M. Vecera, M. Ruckova, T. Kazda, R. Jancalek, M. Hendrych, M. Hermanova, V. Vybihal, P. Fadrus, M. Smrcka, F. Sokol, V. Kubes, R. Lipina, O. Slaby, L. Kren, and J. Sana. 2022. HLA-E and HLA-F Are Overexpressed in Glioblastoma and HLA-E Increased After Exposure to Ionizing Radiation. Cancer Genomics Proteomics 19:151–162.

Jain, N., Z. Zhao, R.P. Koche, C. Antelope, Y. Gozlan, A. Montalbano, D. Brocks, M. Lopez, A. Dobrin, Y. Shi, G. Gunset, T. Giavridis, and M. Sadelain. 2023. Disruption of SUV39H1-mediated H3K9 methylation sustains CAR T cell function. Cancer Discov

Kaiser, B.K., F. Barahmand-Pour, W. Paulsene, S. Medley, D.E. Geraghty, and R.K. Strong. 2005. Interactions between NKG2x immunoreceptors and HLA-E ligands display overlapping affinities and thermodynamics. J Immunol 174:2878–2884.

Kaiser, B.K., J.C. Pizarro, J. Kerns, and R.K. Strong. 2008. Structural basis for CD94/NKG2A recognition of HLA-E. Proc Natl Acad Sci U S A 105:6696–6701.

Kalbasi, A., and A. Ribas. 2020. Tumour-intrinsic resistance to immune checkpoint blockade. Nat Rev Immunol 20:25–39.

Kamiya, T., S.V. Seow, D. Wong, M. Robinson, and D. Campana. 2019. Blocking expression of inhibitory receptor NKG2A overcomes tumor resistance to NK cells. J Clin Invest 129:2094–2106.

Kraehenbuehl, L., C.H. Weng, S. Eghbali, J.D. Wolchok, and T. Merghoub. 2022. Enhancing immunotherapy in cancer by targeting emerging immunomodulatory pathways. Nat Rev Clin Oncol 19:37–50.

Lanier, L.L. 2009. DAP10- and DAP12-associated receptors in innate immunity. Immunol Rev 227:150–160.

Lanier, L.L., B. Corliss, J. Wu, and J.H. Phillips. 1998. Association of DAP12 with activating CD94/NKG2C NK cell receptors. Immunity 8:693–701.

Larson, R.C., M.C. Kann, S.R. Bailey, N.J. Haradhvala, P.M. Llopis, A.A. Bouffard, I. Scarfo, M.B. Leick, K. Grauwet, T.R. Berger, K. Stewart, P.V. Anekal, M. Jan, J. Joung, A. Schmidts, T. Ouspenskaia, T. Law, A. Regev, G. Getz, and M.V. Maus. 2022. CAR T cell killing requires the IFNgammaR pathway in solid but not liquid tumours. Nature 604:563–570.

Le Drean, E., F. Vely, L. Olcese, A. Cambiaggi, S. Guia, G. Krystal, N. Gervois, A. Moretta, F. Jotereau, and E. Vivier. 1998. Inhibition of antigen-induced T cell response and antibody-induced NK cell cytotoxicity by NKG2A: association of NKG2A with SHP-1 and SHP-2 protein-tyrosine phosphatases. Eur J Immunol 28:264–276.

Lee, N., M. Llano, M. Carretero, A. Ishitani, F. Navarro, M. Lopez-Botet, and D.E. Geraghty. 1998. HLA-E is a major ligand for the natural killer inhibitory receptor CD94/NKG2A. Proc Natl Acad Sci U S A 95:5199–5204.

Liu, X., J. Song, H. Zhang, X. Liu, F. Zuo, Y. Zhao, Y. Zhao, X. Yin, X. Guo, X. Wu, H. Zhang, J. Xu, J. Hu, J. Jing, X. Ma, and H. Shi. 2023. Immune checkpoint HLA-E:CD94-NKG2A mediates evasion of circulating tumor cells from NK cell surveillance. Cancer Cell 41:272–287 e279.

Maher, J., R.J. Brentjens, G. Gunset, I. Riviere, and M. Sadelain. 2002. Human T-lymphocyte cytotoxicity and proliferation directed by a single chimeric TCRzeta /CD28 receptor. Nat Biotechnol 20:70–75.

Majzner, R.G., and C.L. Mackall. 2018. Tumor Antigen Escape from CAR T-cell Therapy. Cancer Discov 8:1219–1226.

Malmberg, K.J., V. Levitsky, H. Norell, C.T. de Matos, M. Carlsten, K. Schedvins, H. Rabbani, A. Moretta, K. Soderstrom, J. Levitskaya, and R. Kiessling. 2002. IFN-gamma protects short-term ovarian carcinoma cell lines from CTL lysis via a CD94/NKG2A-dependent mechanism. J Clin Invest 110:1515–1523.

McMahon, C.W., and D.H. Raulet. 2001. Expression and function of NK cell receptors in CD8+ T cells. Curr Opin Immunol 13:465–470.

Miller, J.D., D.A. Weber, C. Ibegbu, J. Pohl, J.D. Altman, and P.E. Jensen. 2003. Analysis of HLA-E peptide-binding specificity and contact residues in bound peptide required for recognition by CD94/NKG2. J Immunol 171:1369–1375.

Mittelbronn, M., P. Simon, C. Loffler, D. Capper, B. Bunz, P. Harter, H. Schlaszus, A. Schleich, G. Tabatabai, B. Goeppert, R. Meyermann, M. Weller, and J. Wischhusen. 2007. Elevated HLA-E levels in human glioblastomas but not in grade I to III astrocytomas correlate with infiltrating CD8+ cells. J Neuroimmunol 189:50–58.

Nguyen, S., V. Beziat, N. Dhedin, M. Kuentz, J.P. Vernant, P. Debre, and V. Vieillard. 2009. HLA-E upregulation on IFN-gamma-activated AML blasts impairs CD94/NKG2A-dependent NK cytolysis after haplo-mismatched hematopoietic SCT. Bone Marrow Transplant 43:693–699.

Riviere, I., K. Brose, and R.C. Mulligan. 1995. Effects of retroviral vector design on expression of human adenosine deaminase in murine bone marrow transplant recipients engrafted with genetically modified cells. Proc Natl Acad Sci U S A 92:6733–6737.

Salome, B., J.P. Sfakianos, D. Ranti, J. Daza, C. Bieber, A. Charap, C. Hammer, R. Banchereau, A.M. Farkas, D.F. Ruan, S. Izadmehr, D. Geanon, G. Kelly, R.M. de Real, B. Lee, K.G. Beaumont, S. Shroff, Y.A. Wang, Y.C. Wang, T.H. Thin, M. Garcia-Barros, E. Hegewisch-Solloa, E.M. Mace, L. Wang, T. O’Donnell, D. Chowell, R. Fernandez-Rodriguez, M. Skobe, N. Taylor, S. Kim-Schulze, R.P. Sebra, D. Palmer, E. Clancy-Thompson, S. Hammond, A.O. Kamphorst, K.J. Malmberg, E. Marcenaro, P. Romero, R. Brody, M. Viard, Y. Yuki, M. Martin, M. Carrington, R. Mehrazin, P. Wiklund, I. Mellman, S. Mariathasan, J. Zhu, M.D. Galsky, N. Bhardwaj, and A. Horowitz. 2022. NKG2A and HLA-E define an alternative immune checkpoint axis in bladder cancer. Cancer Cell 40:1027–1043 e1029.

Sharma, P., S. Goswami, D. Raychaudhuri, B.A. Siddiqui, P. Singh, A. Nagarajan, J. Liu, S.K. Subudhi, C. Poon, K.L. Gant, S.M. Herbrich, S. Anandhan, S. Islam, M. Amit, G. Anandappa, and J.P. Allison. 2023. Immune checkpoint therapy-current perspectives and future directions. Cell 186:1652–1669.

Sharma, P., S. Hu-Lieskovan, J.A. Wargo, and A. Ribas. 2017. Primary, Adaptive, and Acquired Resistance to Cancer Immunotherapy. Cell 168:707–723.

van Montfoort, N., L. Borst, M.J. Korrer, M. Sluijter, K.A. Marijt, S.J. Santegoets, V.J. van Ham, I. Ehsan, P. Charoentong, P. Andre, N. Wagtmann, M.J.P. Welters, Y.J. Kim, S.J. Piersma, S.H. van der Burg, and T. van Hall. 2018. NKG2A Blockade Potentiates CD8 T Cell Immunity Induced by Cancer Vaccines. Cell 175:1744–1755 e1715.

Wischhusen, J., M.A. Friese, M. Mittelbronn, R. Meyermann, and M. Weller. 2005. HLA-E protects glioma cells from NKG2D-mediated immune responses in vitro: implications for immune escape in vivo. J Neuropathol Exp Neurol 64:523–528.

Wolpert, F., P. Roth, K. Lamszus, G. Tabatabai, M. Weller, and G. Eisele. 2012. HLA-E contributes to an immune-inhibitory phenotype of glioblastoma stem-like cells. J Neuroimmunol 250:27–34.

Wu, Z., J. Liang, Z. Wang, A. Li, X. Fan, and T. Jiang. 2020. HLA-E expression in diffuse glioma: relationship with clinicopathological features and patient survival. BMC Neurol 20:59.

