## Supplemental Material for "Targeting HLA-E Positive Cancers with a Novel NKG2A/C Switch Receptor"

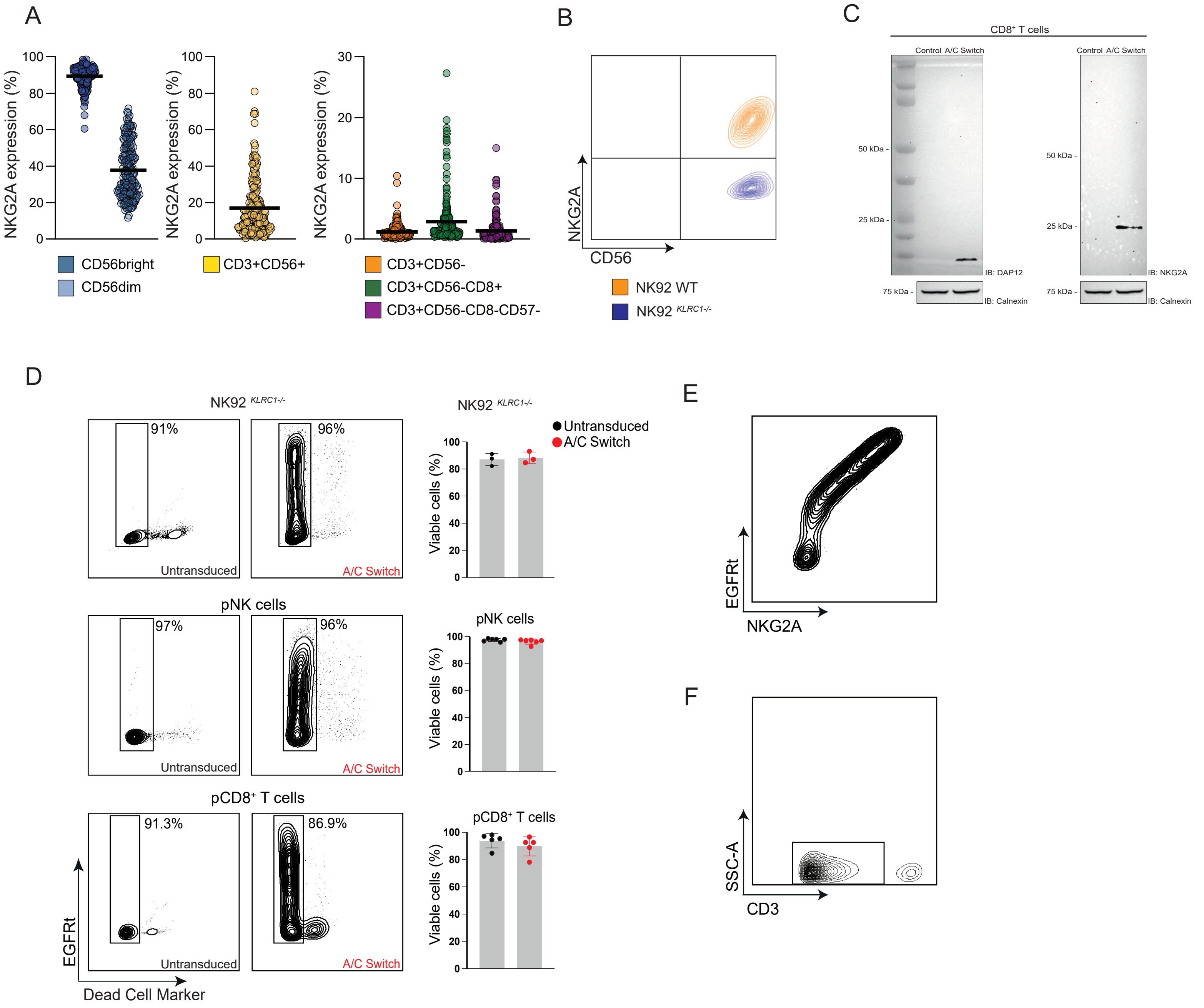
A
NKG2A expression (%)
CD56bright
CD56dim
CD3+CD56+
CD3+CD56-
CD3+CD56-CD8+
CD3+CD56-CD8-CD57-
B
NKG2A
CD56
NK92 WT
NK92 KLRC1-/-
C
CD8+ T cells
Control A/C Switch
50 kDa -
25 kDa -
75 kDa -
IB: DAP12
IB: NKG2A
IB: Calnexin
D
NK92 KLRC1-/-
91%
96%
Untransduced
A/C Switch
Untransduced
A/C Switch
Viable cells (%)
pNK cells
97%
96%
pCD8+ T cells
91.3%
86.9%
EGFRt
Dead Cell Marker
E
EGFRt
NKG2A
F
SSC-A
CD3

Figure S1

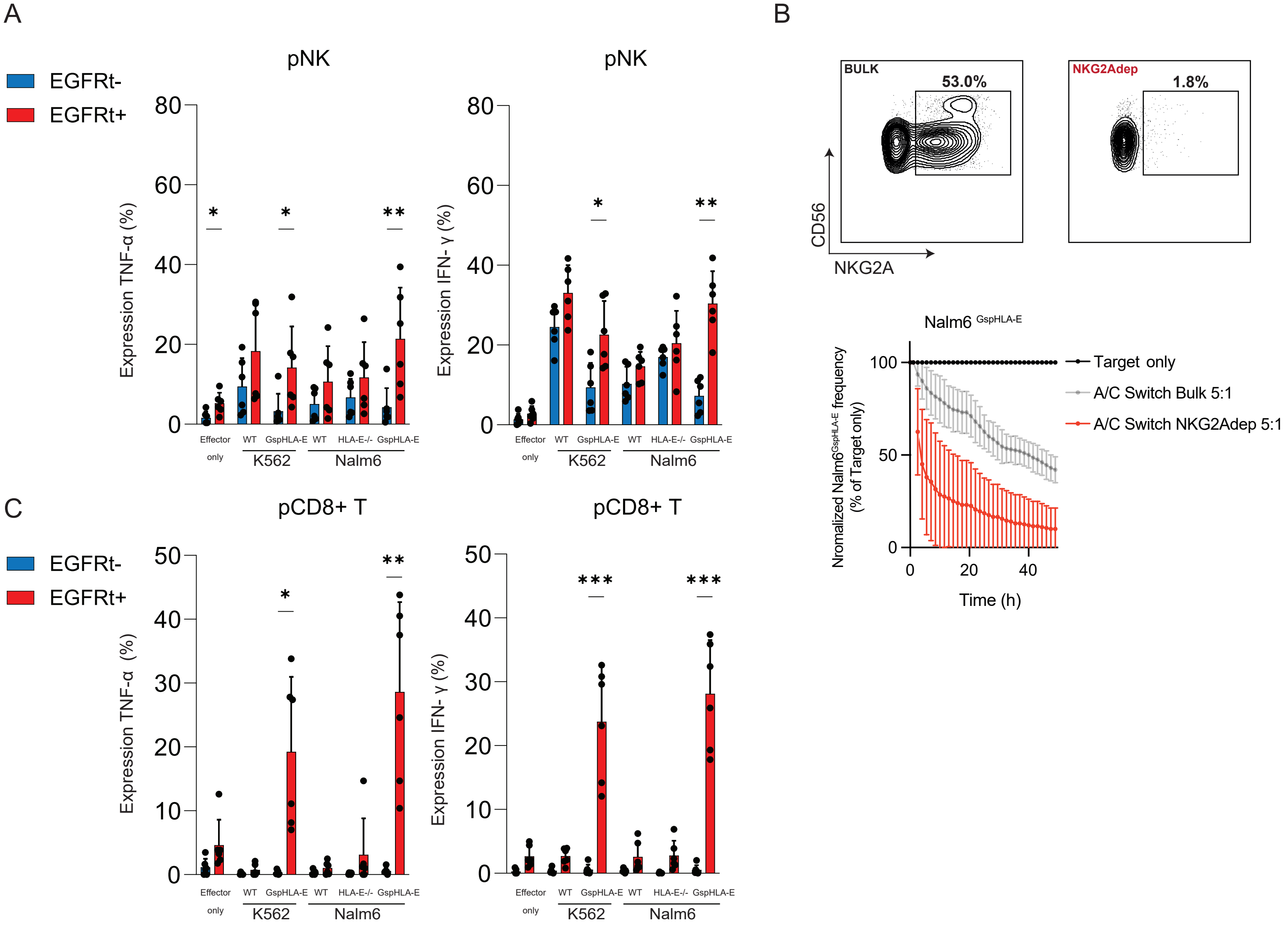
A
pNK
EGFRt-
EGFRt+
Expression TNF-α (%)
Expression IFN- γ (%)
Effector only
WT
GspHLA-E
HLA-E-/-
K562
Nalm6
B
BULK
53.0%
NKG2Adep
1.8%
CD56
NKG2A
Nalm6 GspHLA-E
Nromalized Nalm6GspHLA-E frequency (% of Target only)
Time (h)
Target only
A/C Switch Bulk 5:1
A/C Switch NKG2Adep 5:1
C
pCD8+ T

Figure S2

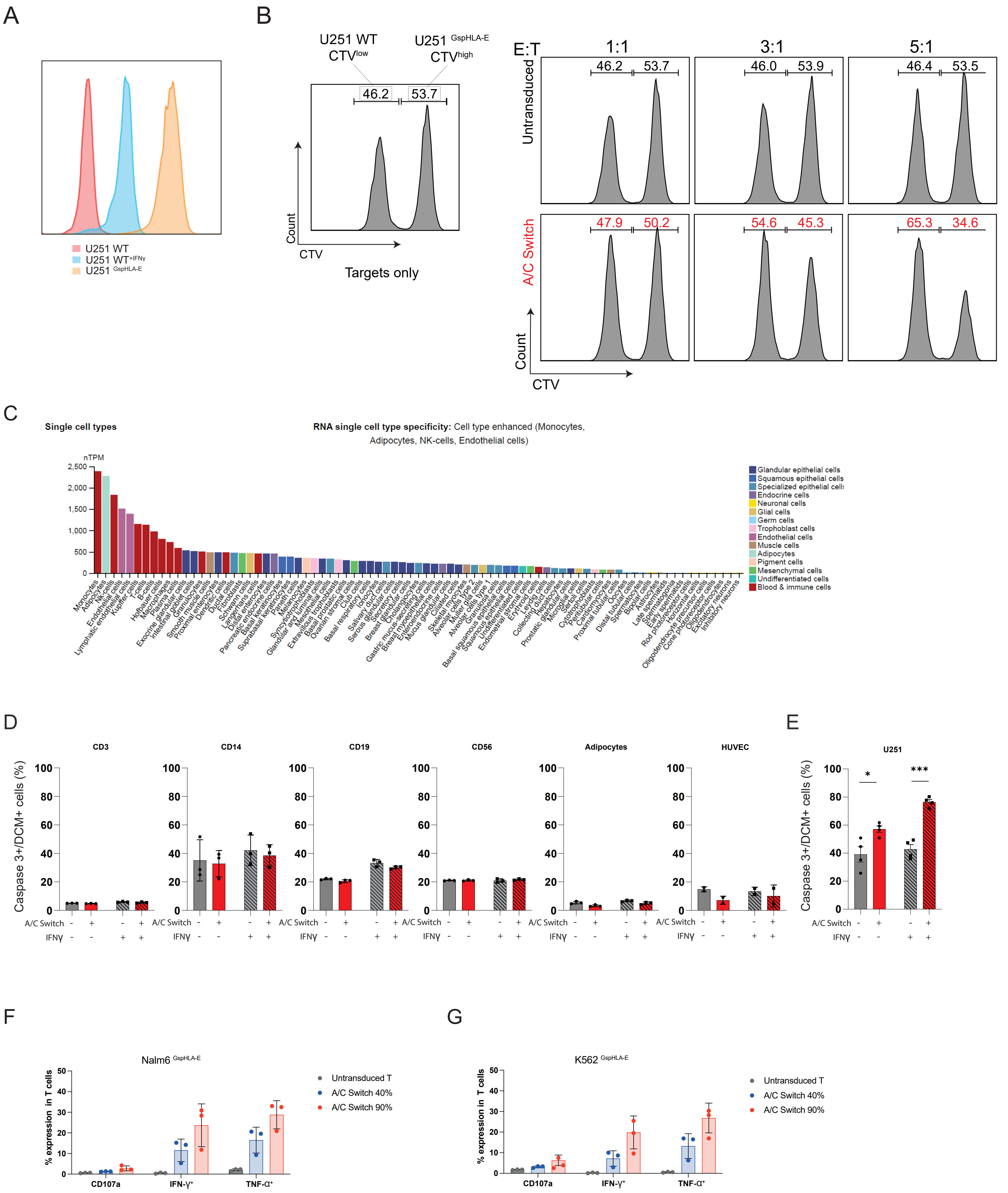
A
U251 WT
U251 WT+IFNγ
U251 GspHLA-E
B
U251 WT CTVlow
U251 GspHLA-E CTVhigh
46.2
53.7
Count
CTV
Targets only
E:T
1:1
3:1
5:1
Untransduced
46.2 53.7
46.0 53.9
46.4 53.5
A/C Switch
47.9 50.2
54.6 45.3
65.3 34.6
Count
CTV
C
Single cell types
RNA single cell type specificity: Cell type enhanced (Monocytes, Adipocytes, NK-cells, Endothelial cells)
nTPM
Glandular epithelial cells
Squamous epithelial cells
Specialized epithelial cells
Endocrine cells
Neuronal cells
Glial cells
Germ cells
Trophoblast cells
Endothelial cells
Muscle cells
Adipocytes
Pigment cells
Mesenchymal cells
Undifferentiated cells
Blood & immune cells
D
CD3
CD14
CD19
CD56
Adipocytes
HUVEC
Caspase 3+/DCM+ cells (%)
A/C Switch
IFNγ
E
U251
Caspase 3+/DCM+ cells (%)
A/C Switch
IFNγ
F
Nalm6 GspHLA-E
% expression in T cells
CD107a
IFN-γ+
TNF-α+
Untransduced T
A/C Switch 40%
A/C Switch 90%
G
K562 GspHLA-E
% expression in T cells
CD107a
IFN-γ+
TNF-α+
Untransduced T
A/C Switch 40%
A/C Switch 90%

Figure S3

**Material and Methods Reagents list**

**Supplementary Table 1**

| **Cells** | **Catalog** | **Provider** |
|---|---|---|
| K562 | CCL-243 | ATCC |
| K562 GspHLA-E | - | *Generated in-house* |
| Nalm6 | CRL-3273 | ATCC |
| Nalm6 HLA-E-/- | - | *Generated in-house* |
| Nalm6 GspHLA-E | - | *Generated in-house* |
| U251 | 9063001 | Millipore Sigma |
| U251 GspHLA-E | - | *Generated in-house* |
| Human adipose-derived stem cells | R7788110 | ThermoFisher |
| Human Umbilical Vein Endothelial Cells, HUVEC | C01510C | ThermoFisher |
| NK92 | CRL-2407 | ATCC |
| NK92KLRC1-/- | - | *Generated in-house* |
| **Flow cytometry antibodies** | **Catalog** | **Provider** |
| CD3-FITC (clone UCHT1)<br>UCHT1)<br>UCHT1 | 300440 | BioLegend |
| CD4-BV785 (clone SK3) | 344642 | BioLegend |
| CD8a-BV711 (clone RPA-T8) | 301044 | BioLegend |
| CD8a-APC (clone RPA-T8) | 301014 | BioLegend |
| CD8a-BUV395 (clone RPA-T8) | 563795 | BD |
| CD14-eFluor450 (clone 61D3) | 48-0149-42 | ThermoFisher |
| CD14-APC (clone M5E2) | 301808 | BioLegend |
| CD19-APC (clone HIB19) | 302212 | BioLegend |
| CD19-V500 (clone HIB19) | 561121 | BD |
| CD25-AF700 (clone BC96) | 302622 | BioLegend |
| CD45RA-APCCy7 (clone HI100) | 304128 | BioLegend |
| CD56-BV711 (clone 5.1H11) | 362542 | BioLegend |
| CD56-ECD (clone N901) | A82943 | Beckman Coulter |
| CD56-BUV395 (clone NCAM16.2) | 563554 | BD |
| CD57-BV605 (clone QA17A04) | 393304 | BioLegend |
| CD57-PE-Dazzle594 (clone HNK-1) | 359620 | BioLegend |
| CD107a-AF488 (clone H4A3) | 328610 | BioLegend |
| HLA-E-APC (clone 3D12) | 342606 | BioLegend |
| NKG2A-APC (clone Z199) | A60797 | Beckman Coulter |
| NKG2A-PECy7 (clone S19004C) | 375114 | BioLegend |

| | | |
|---|---|---|
| NKG2C-PE (REA205) | 130-119-776 | Miltenyi |
| EGFRt-BV421 (clone AY13) | 352911 | BioLegend |
| EGFRt-PE (clone AY13) | 352904 | BioLegend |
| TIM3-BV421 (clone F38-2E2) | 345008 | BioLegend |
| LAG3-BV650 (clone 11C3C65) | 369316 | BioLegend |
| PD-1-BV711 (clone EH12.2H7) | 329928 | BioLegend |
| INFg-APC (clone B27) | 554702 | BD |
| TNFa-PE (clone MAb11) | 554513 | BD |
| TNFa-BV711 (clone MAb11) | 502940 | BioLegend |
| DAP12-PE (REA 900) | 130-115-087 | Miltenyi |
| **Westernblot antibodies** | **Catalog** | **Provider** |
| DAP12 | 12492 | Cell Signaling Tech |
| NKG2A | 10935-1-AP | ProteinTech |
| HLA-E | ab2216 | Abcam |
| Calnexin | NB100-1965 | Bio-Techne |
| Secondary HRP-conjugated anti-mouse | 7076S | Cell Signaling Tech |
| Secondary HRP-conjugated anti-rabbit | 7074S | Cell Signaling Tech |
| **Reagents** | **Catalog** | **Provider** |
| RPMI | 21875-034 | ThermoFisher |
| DMEM | D5796-500ML | Sigma-Aldrich |
| alpha-MEM | 11520616 | ThermoFisher |
| Human Large Vessel Endothelial Cell Basal Medium | M-200-500 | ThermoFisher |
| Fetal Calf Serum (FCS) | F7524-500ml | Sigma-Aldrich |
| Horse Serum | 16050122 | ThermoFisher |
| Penicillin-Streptomycin (PenStrep) | P4333-100ML | Sigma-Aldrich |
| Lymphoprep | 04-03-9391/03 | Serumwerk |
| IFN-g | 285-IF-100 | R&D Systems |
| IL-2 | 200-02-1MG | ThermoFisher |
| IL-7 | 200-07 | Peprotech |
| IL-15 | 130-095-764 | Miltenyi Biotech |
| LIVE/DEAD Near-IR dead cell marker (DCM) dye | L10119 | Invitrogen |
| LIVE/DEAD Aqua dead cell marker (DCM) dye | L34957 | Invitrogen |
| Fixable Viability Dye eFluor 780 | 65-0865-14 | ThermoFisher |
| Flat 6-well plate (non-treated) | 150239 | ThermoFisher |
| Flat 96-well plate | 167008 | ThermoFisher |

| | | |
|---|---|---|
| V-bottomed 96-well plate | 353263 | Falcon |
| RIPA Lysis and Extraction Buffer | 89900 | ThermoFisher |
| Halt™ Inhibitor Single-Use Cocktail (100X) | 78442 | ThermoFisher |
| Skim milk powder | 1.15363.0500 | Sigma-Aldrich |
| NuPAGE™ MOPS SDS Running Buffer (20X) | NP0001 | ThermoFisher |
| NuPAGE™ 4 to 12%, Bis-Tris | NP0321BOX | ThermoFisher |
| NuPAGE™ LDS Sample Buffer (4X) | NP0007 | ThermoFisher |
| NuPAGE™ Sample Reducing Agent (10X) | NP0009 | ThermoFisher |
| SuperSignal™ West Dura Extended Duration Substrate | 34076 | ThermoFisher |
| NK cell isolation kit | 130-092-657 | Miltenyi Biotec |
| CD8+ cell isolation kit | 130-096-495 | Miltenyi Biotec |
| CD4+ cell isolation kit | 130-096-533 | Miltenyi Biotec |
| Pan T cell isolation kit | 130-096-535 | Miltenyi Biotec |
| CD14+ Microbeads | 130-050-201 | Miltenyi Biotec |
| CD19+ Microbeads | 130-050-301 | Miltenyi Biotec |
| NKG2A antibody, Biotin REAfinity | 130-113-564 | Miltenyi Biotec |
| Biotin Microbeads | 130-090-485 | Miltenyi Biotec |
| LS Columns | 130-042-401 | Miltenyi Biotec |
| Retronectin | T100B-TAK | AH diagnostics |
| Anti-CD3/CD28 Dynabeads | 11131D | ThermoFisher |
| Perm/Wash | 51-2091KZ | BD |
| Cytofix/Cytoperm | 51-2090KZ | BD |
| GolgiStop | 51-2092KZ | BD |
| CellTrace Violet Proliferation kit | C34557 | Invitrogen |
| Caspase-3 activity kit (red) | ab65617 | Abcam |
| SF Cell Line 4D-Nucleofector X Kit | V4XC-2012 | Lonza |
| P3 Primary cell 4-D Nucleofector X Kit | V4XP-3012 | Lonza |
| **In vivo supplies** | **Catalog** | **Provider** |
| NSG mice | IMSR_JAX:005557 | Jackson Laboratories |
| Hamilton syringes | 8928B60 | Thomas Scientific |
| Ketamine | 71173 | Covetrus |
| Dexmedetomidine | 72960 | Covetrus |
| Meloxicam | 49756 | Covetrus |
| VetBond Tissue Adhesive | NC2386718 | Fisher Scientific |
| Small Animal Stereotaxic instrument | 900LS | Kopf |
| Ideal Micro Drill Kit | MD-1200 | Braintree Scientific |

| | | |
|---|---|---|
| IVIS Spectrum in vivo imaging system | 124262 | Perkin Elmer |
| D-luciferin | LUCK-1G | Gold Biotechnology |
| **CRISPR/Cas9** | | |
| HLA-E | | Synthego |
| NKG2A | | Synthego |
| Cas9 | | QB3 Berkeley MacroLab |
| TRAC | | Synthego |
